# The biogeographic history of eelpouts and related fishes: linking phylogeny, environmental change, and patterns of dispersal in a globally distributed fish group

**DOI:** 10.1101/2021.01.11.426281

**Authors:** Scott Hotaling, Marek L. Borowiec, Luana S.F. Lins, Thomas Desvignes, Joanna L. Kelley

## Abstract

Modern genetic data sets present unprecedented opportunities to understand the evolutionary origins of taxonomic groups comprising hundreds to thousands of species. When the timing of key events are known, it is also possible to investigate biogeographic history in the context of major phenomena (e.g., continental drift). In this study, we investigated the biogeographic history of the suborder Zoarcoidei, a globally distributed fish group that includes species inhabiting both poles and multiple taxa that produce antifreeze proteins to survive chronic subfreezing temperatures. We first generated a multi-locus, time-calibrated phylogeny for the group. We then used biogeographic modeling to reconstruct ancestral ranges across the tree and quantify the type and frequency of biogeographic events (e.g., founder, dispersal). With these results, we considered how the cooling of the Southern and Arctic Oceans, which reached their present-day subfreezing temperatures 10-15 million years ago (Mya) and 2-3 Mya, respectively, may have shaped the evolutionary history of Zoarcoidei, with an emphasis on the most speciose and widely distributed family, eelpouts (family Zoarcidae). Our phylogenetic results clarified standing issues in the Zoarcoidei taxonomy and showed that the group began to diversify in the Oligocene ∼31-32 Mya, with the center of origin for all families in north temperate waters. Within-area speciation was the most common biogeographic event in the group’s history (80% of all events) followed by dispersal (20%). Finally, we found mixed evidence for polar ocean cooling underpinning Zoarcoidei diversification, with support limited to eelpout speciation in the Southern Ocean over the last 10 million years.

## 1. Introduction

Clarifying spatial origins of diversification and the evolution of geographic ranges is key to understanding patterns of global biodiversity. By considering contemporary distributions in a phylogenetic context, it is possible to assess how key events (e.g., dispersal, extinction, speciation) shape range evolution and diversification (Dupin et al., 2017). With the ever-expanding availability of genetic data in public repositories (e.g., GenBank), declining costs for generating new data, and emerging statistical tools [e.g., biogeographic stochastic mapping (BSM), Matzke (2014)], there has never been a better time to explore complex biogeographic histories across large phylogenies. Cosmopolitan clades, where a single group is distributed throughout all or most of the world, present interesting biogeographical scenarios because no taxonomic group begins with a global distribution and thus many dispersal and vicariance events must occur during its evolution (Nauheimer et al., 2012). Moreover, long-term biogeographic shifts do not occur in an environmentally static landscape. While a group is evolving, diversifying, and shifting its range over millennia, the habitats it occupies are also changing in both size and suitability. Large-scale environmental shifts can drive species’ radiations and when the timing of influential events (e.g., the separation of two land masses or cooling of a major ocean) are known, then it is possible to test hypotheses linking biogeographic patterns to processes on a calibrated timeline (Dupin et al., 2017).

A cosmopolitan group of particular biogeographical interest are eelpouts (family Zoarcidae), the most speciose family in the suborder Zoarcoidei, comprising ∼75% of the suborder’s ∼400 species (Fricke et al., 2018), and representing the only Zoarcoidei family with species that inhabit both poles (Møller et al., 2005). Eelpouts are also one of the most rapidly speciating fish clades, with their propensity for deep-waters and high-latitudes implicated as potential drivers of their high speciation rate (Rabosky et al., 2018). At polar latitudes, marine environments are chronically cold, and often subfreezing, yet they retain high levels of biological productivity and species richness (DeVries and Steffensen, 2005). Considerable focus has been devoted to understanding how and when organisms diversified in the Southern and Arctic Oceans (e.g., González-Wevar et al., 2010; Hopkins and Marincovich Jr, 1984), particularly as it relates to when both oceans reached their contemporary subfreezing temperatures [Southern Ocean: 10-15 million years ago (Mya), Arctic Ocean: 2-3 Mya; DeVries and Steffensen (2005)]. Generally speaking, most Zoarcoidei species are found in the Northern Hemisphere, specifically the northwestern Pacific Ocean, which has been proposed as a speciation center for the group (Anderson, 1994; Shmidt, 1950).

A key innovation among the Zoarcoidei is the evolution of antifreeze proteins (AFP). AFPs have evolved repeatedly across the Tree of Life, including in multiple fish lineages beyond the Zoarcoidei (e.g., Antarctic notothenioids, Chen et al., 1997) and have been hypothesized to be a major factor underlying adaptive radiations in some groups (e.g., notothenioids, Matschiner et al., 2011). Adaptive radiations occur when high speciation rates, common ancestry, and a phenotype-environment correlation drive a rapid increase in species diversity and often stem from ecological opportunity (Schluter, 2000). For instance, the Antarctic notothenioid adaptive radiation into freezing Antarctic waters has been linked, in part, to the evolution of AFPs (Matschiner et al., 2011; Near et al., 2012). Within the Zoarcoidei, AFPs are present in at least five families—Anarhichadidae, Cryptacanthodidae, Pholidae, Stichaeidae, Zoarcidae (Davies et al., 2002; Davies et al., 1988)—with AFP-containing lineages inhabiting Arctic and Antarctic waters. Thus, the contemporary distributions of Zoarcoidei species, and particularly eelpouts living at both poles with their associated AFPs, raise questions about how cooling of the Arctic and Southern Oceans may have influenced the group’s evolutionary history.

Here, we used multi-locus sequence data to construct a time-calibrated, comprehensive phylogeny of the suborder Zoarcoidei. Next, we used this phylogeny to clarify issues of taxonomic uncertainty in the group and better understand its biogeographic history. To the first, previous phylogenetic efforts have noted issues with the Zoarcoidei taxonomy, primarily stemming from a lack of monophyly in the Stichaeidae family, which led to the description of two new families, Eulophiidae and Neozarcidae (Kwun and Kim, 2013). We confirm and build upon these prior efforts to improve Zoarcoidei taxonomy. To the second—biogeographic history—we reconstructed ancestral ranges for every node of our phylogeny and considered what, if any, evidence exists for cooling of the Arctic and Southern Oceans to have driven patterns of speciation. We performed biogeographic stochastic mapping on our phylogeny to quantify the types of biogeographic events (e.g., founder-event speciation, dispersal) that have underpinned the group’s diversification. To our specific question of whether ocean cooling has been a major driver of speciation within Zoarcoidei, and for eelpouts in particular since they are the only globally distributed family in the suborder, we expected to observe three lines of evidence: (1) higher support for biogeographic models that incorporate Arctic and Southern Ocean cooling, (2) bursts of speciation following the cooling of each ocean at roughly 10 (Southern) and 2 (Arctic) Mya, and (3) more dispersal events into the Arctic and Antarctic than out of them as cold-adapted Zoarcoidei took advantage of new ecological opportunity.

## 2. Materials and Methods

### 2.1. Data collection

We obtained sequence data for up to three nuclear genes (*rag1, rho, rnf213*) and three mitochondrial genes [*cytochrome oxidase I* (*mt-co1*), *cytochrome B* (*mt-cyb*), *16S rRNA* (*16S*)] from 223 specimens in the suborder Zoarcoidei and an outgroup, *Eleginops maclovinus* (suborder Notothenioidei). Our data set included a combination of existing data in GenBank and newly generated data (Table S1). For phylogenetic biogeographic modeling and ancestral range reconstruction (see *2.3 Biogeographic modeling and ancestral range estimation*), it was important that we binned species’ contemporary distributions into geographic categories. We first defined the geographic distribution of each species in our data set using FishBase (http://fishbase.org; Froese and Pauly, 2019), an online database with species-level distribution information that stems from published literature and observations reported on the Ocean Biogeographic Information System (OBIS, https://obis.org/; Grassle, 2000) and the Global Biodiversity Information Facility (GBIF, https://www.gbif.org/; Lane and Edwards, 2007). We then binned contemporary distributions for each species into five geographic zones based on their latitudinal range with multiple zones allowed for a given taxon: (1) Arctic (north of the Arctic Circle, >66.5°N), (2) north temperate (23.5°N − 66.5°N), (3) tropical (between the Tropic of Cancer in the northern hemisphere and the Tropic of Capricorn in the southern hemisphere; 23.5°N to 23.5°S), (4) south temperate (23.5°S − 66.5°S), and (5) Antarctic (south of the Antarctic Circle, >66.5°S).

We collected new sequence data for four species that were field-identified as *Ophthalmolycus amberensis, Lycenchelys tristichodon, Lycodapus endemoscotus*, and *Melanostigma* sp. Using polymerase chain reaction (PCR) and targeted Sanger sequencing. For each taxon, DNA was extracted from frozen tissue (either muscle, liver, or a fin clip) using a MagAttract HMW DNA Kit (Qiagen), following the manufacturer’s protocol for 25 mg tissue samples. We amplified our six markers using primers listed in Table S2 with the same PCR conditions: initial denaturation for 4 min at 94°C, 35 cycles of 30 s at 94°C, 30 s at 55°C and 45 s at 72°C, and a final elongation for 7 min at 72 °C.

We also extracted sequences for *Lycodichthys dearborni* (*rag1, rho, rnf213, mt-co1*, and *mt-cyb*) and *Lycodes polaris* (*rag1, rho, rnf213, mt-cyb*, and *16S*) from short-read genome assemblies. Genomes were assembled from high-coverage (>50x), short-read sequence data (either 100-bp or 150-bp paired-end Illumina sequence data) with SPAdes v3.11.1 and default settings (Bankevich et al., 2012). To extract sequences, we used BLAST+ v2.5.0 (Altschul et al., 1990) to align our primers against each assembly. Matches with an e-value less than 0.5 that were also the longest match between the query and target were identified as our best hits. We extracted the sequence between primers (the target) with bedtools (Quinlan and Hall, 2010). To confirm the identity of sequences, we used BLAST to compare the extracted markers against the NCBI database to verify they were similar to sequences from closely related species.

### 2.2. Phylogenetic reconstruction and divergence timing

Nucleotide sequences for *rag1, rho, rnf213, mt-co1*, and *mt-cytb* were translated to amino acid sequences and aligned using MUSCLE v3.8.31 with default settings (Edgar, 2004). Nucleotide alignments were then generated using the amino acid alignments with PAL2NAL v14-0 (Suyama et al., 2006). Nucleotide sequences for *16S* were aligned using MUSCLE v3.8.31 with default settings (Edgar, 2004). After concatenation, we used the aligned nucleotide data set to estimate phylogeny using maximum likelihood and infer divergence times in a Bayesian framework. To infer the maximum likelihood tree we used IQ-TREE v1.6.10 (Nguyen et al., 2015). We provided partitions based on codon positions in each of the five coding genes and let each partition have an individual rate while sharing branch lengths across partitions (Chernomor et al., 2016). We let IQ-TREE find the best substitution models and partitioning scheme (Kalyaanamoorthy et al., 2017). To improve the thoroughness of the tree search algorithm we decreased the perturbation parameter to 0.3 from a default of 0.5 and increased unsuccessful tree search iterations to 500 from a default of 100. We assessed confidence across the tree with 5,000 replicates of ultrafast bootstrap approximation (Hoang et al., 2018).

We estimated divergence timing under a fossilized birth-death process (Heath et al., 2014) as implemented in MrBayes v3.2.7a (Ronquist et al., 2012). We used the fossil *Proeleginops grandeastmanorum* (family Eleginopsidae, age 38-45 Mya) constrained as sister to the outgroup species *Eleginops maclovinus* (Bienkowska-Wasiluk et al., 2013). Because of uncertainty of their placement, two fossil species—*Agnevicthys gretchinae* and *Palaeopholis laevis* (family Pholidae, age 11.5-12.3 Mya; Nazarkin, 2002)—were allowed to be placed as either the stem (outside of the clade formed by extant species) or crown (within the clade of extant species) for the group during exploration of the tree space. We included several fossils identified as Stichaeidae but because preliminary analysis demonstrated polyphyly of this family, we allowed these fossils to be placed anywhere within the in-group excluding Bathymasteridae: *Nivchia makushoki, Stichaeus brachigrammus*, and *Stichaeopsis sakhalinensis* (age 11.5-12.3 Mya; Nazarkin, 1998), undescribed fossils NSM PV 22683 (age 13-16 Mya) and PIN 3181/1050 (11.6-13.5 Mya; Nazarkin and Yabumoto, 2015), and *Stichaeus matsubarai* (age 5.3-23 Mya; Yabumoto and Uyeno, 1994). We used fossils assigned to the contemporary species *Lycodes pacificus* (family Zoarcidae) to date its age at 0.78-2.59 Mya (Fitch, 1967).

For each fossil, we sampled age from a uniform distribution spanning its possible age range. Because recent work suggests gene-partitioning for divergence dating may result in unrealistically narrow confidence intervals (Angelis et al., 2018), we used an unpartitioned GTR model with gamma rate distribution broken into six discrete categories, the independent gamma-rate relaxed clock model, and extant sample proportion of 0.5. We set the root age prior to be an exponential distribution offset at 38 Mya (the youngest likely age of *P. grandeastmanorum*) with a mean of 70 Mya. We performed these analyses under two scenarios: one assuming taxon sampling was random and one assuming taxon sampling was done to maximize taxonomic diversity (Zhang et al., 2016). The choice of sampling scheme assumption can impact dating analyses if significant mismatch between assumed and actual taxon sampling exists. For example, when only a few species are sampled to represent genera or families in a clade containing thousands of species unequally distributed across these taxa, the sampling scheme is maximizing taxonomic and phylogenetic diversity and is different from a random sample of species from that clade. This can lead to bias in fossilized birth-death process dating (Zhang et al., 2016). For each MrBayes analysis we ran four replicates, each with four chains, for 400 million generations, sampling every 10,000 generations and discarding the first 20% of samples as burn-in. We assessed the reliability of these analyses by confirming that effective sample size for each parameter was greater than 100, potential scale reduction factor values were close to 1.0, proposal acceptance rates were between 20-70%, average standard deviations of split frequencies were below 0.01, and that time-series of parameter values converged across replicates. We did not observe differences between random and diversified sampling. Thus, we used diversified sampling results for downstream analyses. To visualize summarize our results, we generated a lineages through time plot for the full species tree with the *ltt* function of Phytools (Revell, 2012) and plotting in ape (Paradis and Schliep, 2019).

### 2.3. Biogeographic modeling and ancestral range estimation

For biogeographical modeling, we used “BioGeography with Bayesian (and likelihood) Evolutionary Analysis in R Scripts” v1.1.2 (BioGeoBEARS; Matzke, 2014). To identify the best-fit model, we compared likelihoods of six models for ancestral range estimation including dispersal-extinction cladogenesis (DEC; Ree, 2005; Ree and Smith, 2008), dispersal-vicariance analysis (DIVALIKE; Ronquist, 1997), and Bayesian inference of ancestral areas (BAYAREALIKE; Landis et al., 2013), as well as a variant of each model allowing for founder-event speciation (“+*j*” parameter designation). In addition to *j*, the models included two other free parameters: *d* (rate of range expansion) and *e* (rate of range contraction). Because our dated Bayesian consensus tree contained several polytomies, we ran BioGeoBEARS model selection separately on ten randomly chosen posterior trees to account for uncertainty. For all trees, we removed fossil taxa and taxonomic replicates to ensure that each species was represented only once. We also removed tips that were not reliably assigned to a described species (e.g., to genus only) and/or had no sampling locality information given and thus no geographic context.

After binning species into geographic zones as described above—Arctic, North Temperate, Tropical, South Temperate, Antarctic—we ran two types of BioGeoBEARS analyses. (1) “Unconstrained”, meaning dispersal probabilities were equal across time and space and taxa were allowed to have discontinuous ranges (e.g., Arctic and Tropical but not North Temperate).

(2) A more parameter-rich and biologically realistic “time-stratified” analysis with dispersal probabilities modified for three pre-defined time periods—0-3 Mya, 3-20 Mya, and 20 Mya and older (i.e., the time before during and after cooling of the Arctic and Southern Oceans, DeVries and Steffensen, 2005)—to incorporate predicted geographic and ecological distances among range categories. Dispersal was penalized by distance only for the time period before the Southern or Arctic Oceans began cooling (>20 Mya), a dispersal penalty was added for the Antarctic zone after the Southern Ocean began cooling and reached its present state (3-20 Mya), and a dispersal penalty was added for the Arctic zone after the Arctic Ocean began cooling to its present-day temperature (0-3 Mya). For both sets of analyses, a maximum occupancy of three geographic zones was allowed and for the time-stratified analyses, only adjacent ranges were allowed (e.g., Tropical-North Temperate-Arctic). The dispersal matrices used in these analyses are provided in Table S3.

### 2.4. Biogeographic stochastic mapping

In order to quantify the number of each type of biogeographic events in Zoarcoidei evolution we used biogeographic stochastic mapping (Dupin et al., 2017). Six types of biogeographic events were allowed in the models tested: speciation within-area (both species occupy the same area post-speciation), speciation within-area subset (one species inhabits a subset of the range post-speciation), vicariance, founder event, range expansion, and range contraction (see complete descriptions in Dupin et al., 2017). We differentiated among models using the Akaike information criterion corrected for small sample sizes (AICc; Cavanaugh, 1997). According to AICc, “BAYAREALIKE+J” was favored across all ten randomly selected posterior trees for both unconstrained and time-stratified analyses (see *3. Results*). We therefore used BAYAREALIKE+J under the time-stratified regime for biogeographic stochastic mapping with 100 stochastic replicate maps performed on each of the ten randomly chosen posterior trees.

To obtain consensus results we averaged event counts from each of the 10 posterior trees for the best-fit model (BAYAREALIKE+J).

## 3. Results

### 3.1. Data collection

We acquired sequence data for 223 specimens representing at least 198 described species or subspecies from 10 families within Zoarcoidei. This translates to ∼49% of described species diversity (*n* = 403) in the suborder (FishBase; Froese and Pauly, 2019). For three families, we sampled 100% of described diversity: Anarhichadidae, Ptilichthyidae, and Zaproridae. For the most speciose family in the suborder—eelpouts (Zoarcidae)—we sampled 113 of 303 described species (37.3%; Figure 1). Across all specimens and markers, our data set was 44.9% complete with only seven (3.1%) specimens represented by a single marker. Sampled taxa spanned 274 contemporary geographic zones with 42 species in the Arctic (15.1%), 180 species in the North Temperate zone (64.5%), 15 species in the Tropical zone (5.4%), 26 species in the South Temperate zone (9.3%), and 11 species in the Antarctic (3.9%; Table S1). Only eelpouts (Zoarcidae) had distributions in the South Temperate and Antarctic zones (Table S1).

**Figure 1.**
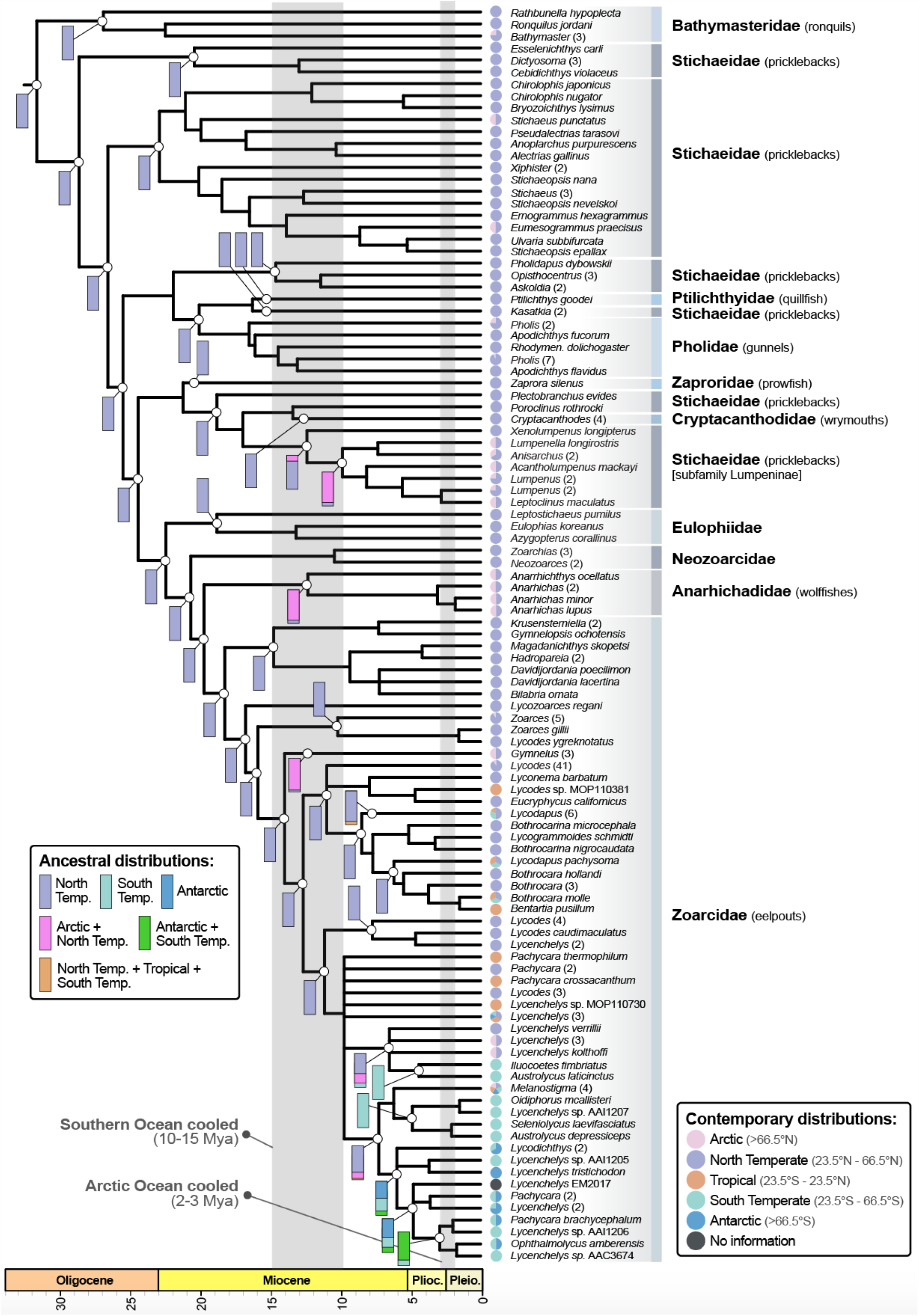
A time-calibrated tree of the suborder Zoarcoidei. For visualization, when multiple species within the same genus formed a monophyletic group, we compressed the group. The number of taxa that were compressed are given in parentheses after the tip label. To the left of nodes, colored areas within vertical rectangles indicate the amount of support for that ancestral distribution group (Note: Up to three geographic zones could be combined for the ancestral range reconstruction). More area indicates more support for that ancestral distribution over others (if applicable). To the right of tips, small pie charts represent present-day distributions across our five latitudinally defined geographic zones (Arctic, North Temperate, Tropical, South Temperate, Antarctic). When multiple tips are compressed into one pie chart and/or a taxon’s range spans multiple regions, the proportion for each region is reflected in the pie chart. Like historical distributions, contemporary distributions were also allowed to span more than one geographic zone. Thus, the number of pie chart components does not necessarily equal the number of taxa in a given group. The tree was rooted with *Eleginops maclovinus* which was removed for visualization. The numeric scale at the bottom of the figure indicates millions of years before present with corresponding geological epochs. Vertical gray bars indicate timing of the cooling of the Southern and Arctic Oceans, respectively. Complete trees (with outgroups) including dating estimates, probabilities for each node, and the full maximum likelihood tree are included in the Supplementary Materials as Figures S1, S2, and S3, respectively.

### 3.2. Phylogenetic reconstruction

Our phylogeny indicates that the Zoarcoidei diverged from the last common ancestor of notothenioids and Zoarcoidei during the Lower Cretaceous period, ∼104 Mya [95% highest posterior density (HPD): 72-152 Mya] and began to radiate in the Oligocene, ∼31-32 Mya (Figures 1, S1). Major families were recovered as monophyletic except for the Stichaeidae which were recovered as polyphyletic, in line with previous studies (e.g., Clardy, 2014; Radchenko, 2016). Our results lend support to the current taxonomy of Eulophiidae and Neozoarcidae which were described by Kwun and Kim (2013) and expanded by Radchenko (2015). We also found support for the genus *Kasatkia* (currently in the Stichaeidae family) as sister to *Ptilichthys goodei*, the only described species in the family Ptilichthyidae (Figure 1). From a timing perspective, the eelpouts (Zoarcidae), the only family with a global distribution, emerged in the early Miocene (∼18 Mya) and have steadily diversified until the present, with only one potential burst of speciation: the largest polytomy in our tree, suggesting rapid speciation, occurred ∼10 Mya when the Southern Ocean had largely cooled to present-day temperatures (Figures 1-2).

**Figure 2.**
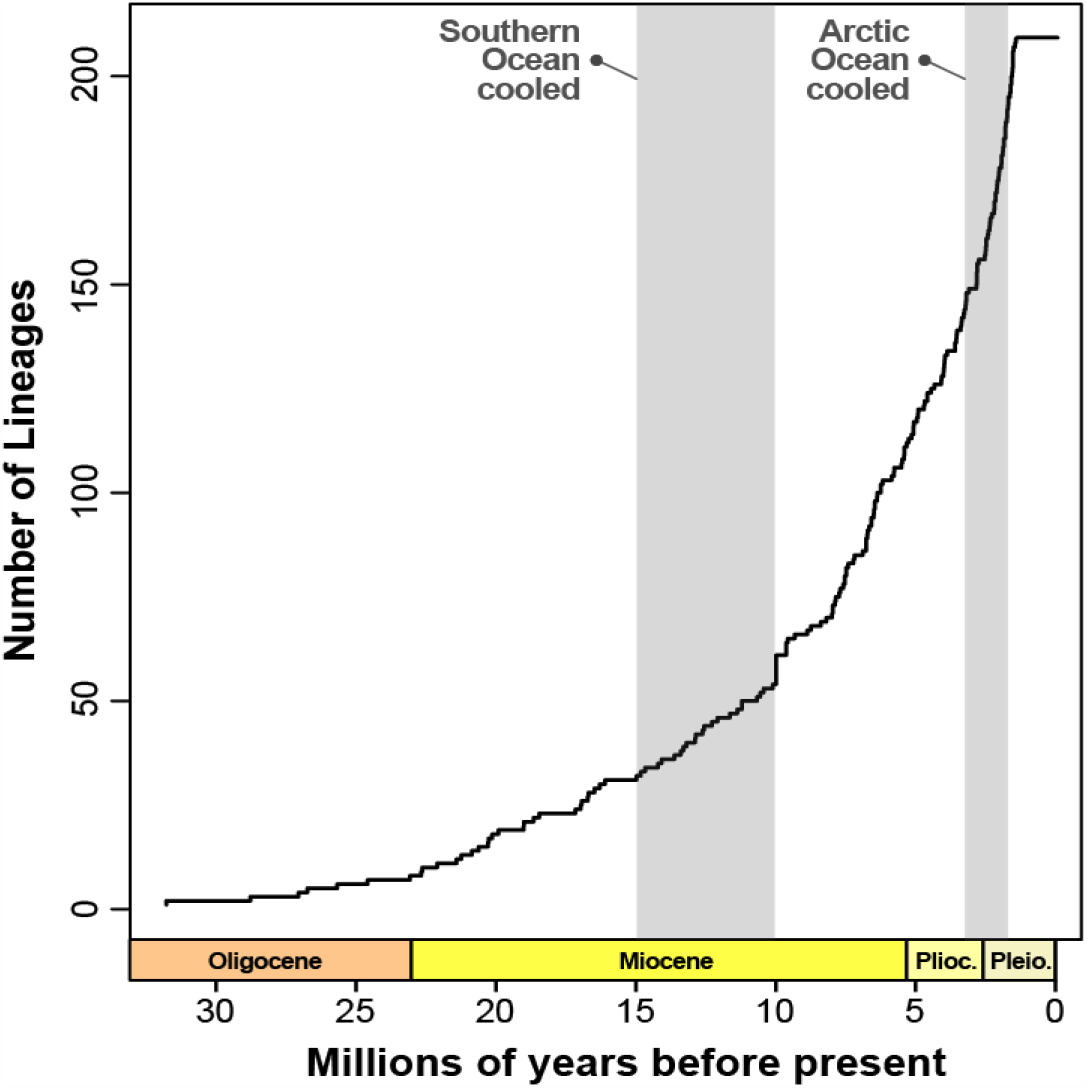
A lineage through time plot for the suborder Zoarcoidei with the timing of Southern and Arctic Ocean cooling noted.

### 3.3. Biogeographic modeling and ancestral range estimation

For both time-stratified and unconstrained analyses, our model selection results strongly favored the BAYAREALIKE +*j* model with the second-best model (DEC+*j*) 75 AICc units higher in both cases (Table 1). In line with similar biogeographic studies on cosmopolitan species (e.g., Dupin et al. 2017), the inclusion of a founder-event speciation parameter (+*j*) substantially improved fit across all models tested (Tables 1, S4). Our time-stratified analyses were also a better fit to the data with a 31 AICc unit difference between the best-fit model (BAYAREALIKE +*j*) for time-stratified versus unconstrained analyses (Table S4). Given this, we focus hereafter on the time-stratified results. Ancestral range reconstruction under the best-fit model (BAYAREALIKE +*j*) supported a North Temperate origin for the entire suborder, as well as every family within the group with the exception of the wolffishes (family Anarhichadidae) with the bulk of support (>80%) in favor of a combined Arctic+North Temperate ancestral range for that group (Figure 1). Two other clades, one including Stichaeidae lineages with four *Lumpenus* species and the other containing three *Gymnelus* eelpout species, also exhibited strong support for an

**Table 1.** A summary of biogeographic model selection for the time-stratified analyses averaged across 10 randomly selected posterior trees to account for polytomies in the consensus tree. Complete model selection results, including those for the “unconstrained” analyses which closely align with those presented here, are included in Table S4. The models tested follow those outlined in (Matzke, 2013) and include dispersal-extinction cladogenesis (DEC; Ree, 2005; Ree and Smith, 2008), dispersal-vicariance analysis (DIVALIKE; Ronquist, 1997), and Bayesian inference of ancestral areas (BAYAREALIKE; Landis et al., 2013) as well as a variant of each allowing for founder-event speciation (+*j*).

| Model | Parameters | Mean AICc | ΔAICc | Model choice |
| --- | --- | --- | --- | --- |
| DEC | 2 | 669.2 | 99.2 | 3 |
| DEC+j | 3 | 645.2 | 75.2 | 2 |
| DIVALIKE | 2 | 721.5 | 151.4 | 6 |
| DIVALIKE+j | 3 | 684.8 | 114.8 | 5 |
| BAYAREALIKE | 2 | 676.8 | 106.8 | 4 |
| BAYAREALIKE+j | 3 | 570.1 | -- | 1 |

Arctic+North Temperate origin. The only non-Northern Hemisphere ancestral range we found support for was within eelpouts, specifically a number of lineages in the subfamily Lycodinae. For example, for a clade containing several *Lycenchelys* and four *Lycodichthys* species, including *Lycodicthys dearborni*, an Antarctic resident known from 72°-78°S, we found ∼50% support for an Antarctic ancestral range followed by ∼40% support for South Temperate, and 10% support for a combination of Antarctic+South Temperate (Figure 1).

### 3.4. Biogeographic stochastic mapping

Across the Zoarcoidei, most biogeographic events were within-area speciation (80%) followed by two types of dispersals: range expansions (13.5%) and founder events (6.5%; Table 2). The fact that we observed a high number of within-area speciation events is unsurprising given that we divided the Earth into five large geographic zones. Similarly, a lack of vicariance events likely reflects the continuous nature of the marine environment with few strong dispersal barriers.

**Table 2.** Summary of biogeographic stochastic mapping results for the suborder Zoarcoidei and the best-fit model (BAYAREALIKE+*j*). The six types of biogeographic events allowed in the model are described fully in Dupin et al. (2017). Speciation within-area and speciation within-area subset differ in that under the former, ranges before and after divergence are the same whereas in the latter, one of the new lineages only occupies a subset of its former range. Included values are averaged [with standard deviations (SD) for the means] across 10 randomly selected posterior trees to account for polytomies in the consensus tree.

| Mode | Type | Mean (SD) | Percent |
| --- | --- | --- | --- |
| Within-area speciation | Speciation within-area | 194.1 (1.4) | 80 |
|  | Speciation within-area subset | 0 | 0 |
| Dispersal | Founder event | 15.9 (1.4) | 6.5 |
|  | Range expansions | 32.7 (2.4) | 13.5 |
|  | Range contractions | 0 | 0 |
| Vicariance | Vicariance | 0 | 0 |
| Total |  | 241.7 (2.4) | 100 |

For dispersal events (i.e., range expansions and founder events), 30% of all events were out of the North Temperate zone with the bulk going into the adjacent Arctic (mean = 13.44 events) or Tropical (9.69) zones (Figure 3A). In general, far fewer dispersal events occurred in the Southern Hemisphere, likely reflecting how much more common Zoarcoidei species are in the Northern Hemisphere, and the North Temperate zone in particular (Figure 1). Range expansion events largely mirrored total dispersal events, with the bulk occurring from North Temperate into the Arctic zone (13.20; Figure 3B). Founder events, however, followed a slightly different pattern with most events occurring from the North Temperate into the Tropical (4.51) and South Temperate (3.47) zones, respectively (Figure 3C). Again, this pattern likely reflects the concentration of Zoarcoidei species in the North Temperate zone (Figure 1).

**Figure 3.**
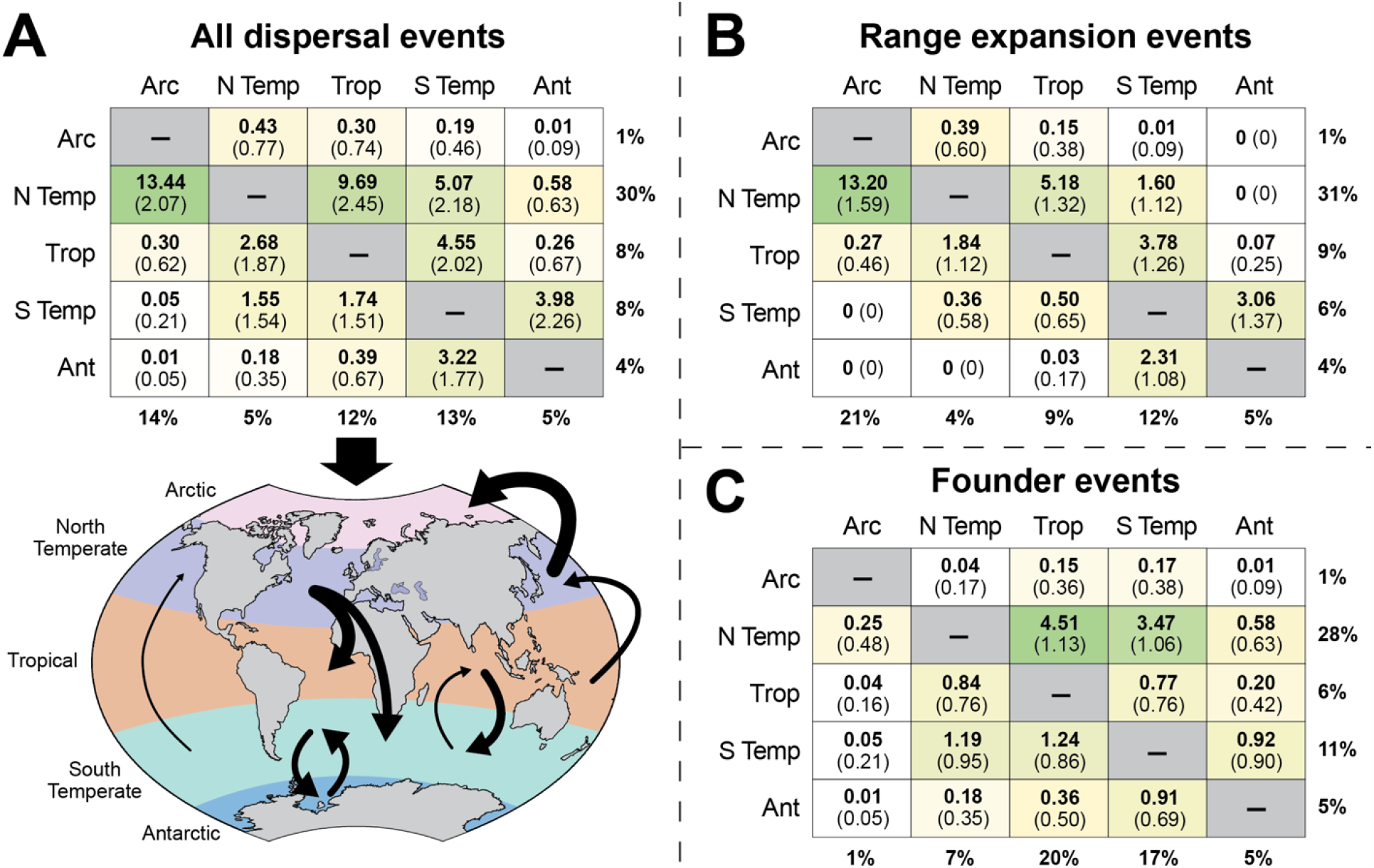
Summary of dispersal events in the history of the Zoarcoidei as estimated with biogeographic stochastic mapping (BSM). Counts of dispersal events (bold) and standard deviations (in parentheses) were averaged across 50 replicate BSMs for each of 10 phylogenies that were randomly sampled from the posterior distribution. (A) Total dispersal events are given in the matrix and are depicted on a global map with colors representing defined geographic zones. Black arrows between areas indicate the frequency and direction of dispersal events. Only events with total mean counts of 1 or more are shown. For visualization, arrow thickness corresponds to the log_10_ of the event count multiplied by 2. Arrows only correspond to individual geographic zones and do not correspond to specific oceans or regions. Their placements within zones are purely for visualization. Total event counts in (A) are divided into the two non-zero types of dispersal events observed in this study in (B) and (C). In (B) and (C), summarizing percentages were calculated for each group separately so cannot be compared between them. Total event counts, however, can be directly compared and sum to the values in (A). Within matrices, color indicates event frequency with darker green shading indicating higher frequencies. Given the counts and associated standard deviations, lower frequency counts (e.g., less than 1) are not necessarily different from zero. For each matrix, rows represent ancestral states where the lineage dispersed from and columns represent descendant states where the lineage dispersed to. The percentage of total events that a row or column comprises in a given matrix are shown in bold font on the margins. Geographic zone abbreviations include Arctic (Arc), North Temperate (N Temp), Tropical (Trop), South Temperate (S Temp), and Antarctic (Ant).

Focusing on the Arctic and Antarctic zones which cooled to their present-day subfreezing temperatures over the last ∼20 Mya, we observed asymmetric dispersal rates for both. Indeed, just 1% of all dispersal events originated from the Arctic whereas it received 14% of all dispersals. The Antarctic zone was less skewed but still showed a slight bias with 4% of all dispersal events originating from it while receiving 5% (Figure 3). Collectively, most of the asymmetry we observed was driven by range expansions into and out of the Arctic; the Arctic received 21% of all range expansions while generating just 1% from within.

## 4. Discussion

Our phylogenetic and biogeographic analyses confirmed that the suborder Zoarcoidei primarily evolved in northern temperate waters (23.5°-66.5°N). This general pattern is true for all families with one exception—the eelpouts (family Zoarcidae)—which exhibit a global distribution with a portion of their species diversity occurring from the Tropics to the Southern Ocean (Figure 1).

Our best-fit biogeographic model included time-stratified matrices that reflected the elevated dispersal challenges of the Arctic and Southern Oceans as they cooled to their contemporary subfreezing temperatures. Support for these time-stratified analyses over models without time-stratification suggests that cooling of both oceans is important to understanding dispersal among the Zoarcoidei. We also observed a clear skew in dispersal directionality during the group’s evolutionary history with both range expansion and founder events much more likely to originate from the North Temperate zone than anywhere else. Finally, we confirmed standing issues with the Zoarcoidei phylogeny, namely a lack of monophyly for Stichaeidae, and we make recommendations to improve these issues below.

### 4.1. Phylogenetic reconstruction and biogeography

Our analyses support diversification of families within the Zoarcoidei occurring ∼31-32 Mya during the Oligocene, beginning with the separation of ronquils (family Bathymasteridae) from the rest of the group. This timing differs from two previous estimates but is closer to the ∼37 Mya estimate from Betancur-R et al. (2013) than the ∼18 Mya estimate of Radchenko (2016), despite using the same markers as Radchenko (2016). In general, all divergences in our reconstruction were deeper in time than those of Radchenko (2016). Betancur-R et al. (2013) included many more taxa and calibrations than Radchenko (2016) and our data set included roughly three times as many specimens.

From an ecological standpoint, the difference between the timing of eelpout (family Zoarcidae) emergence between our study (∼18 Mya) versus the ∼11-13 Mya reported by Radchenko (2016) is important as it places the group’s initial divergence on either side of when the Southern Ocean reached its present-day subfreezing temperature 10-15 Mya (Figure 1). Eelpouts, as well as other high-latitude fish clades (e.g., Antarctic notothens), are one of the fastest speciating fish groups (Rabosky et al., 2018). Thus, it is possible that the cooling of the polar seas, paired with key innovations like the evolution of AFPs (Deng et al., 2010), provided the necessary ecological opportunity and physiological tools necessary for two bursts of eelpout speciation as the Southern and Arctic Oceans cooled. We found some, albeit limited, evidence for this among southern lineages, with a multi-tip polytomy at 10 Mya, soon after the Southern Ocean reached its contemporary subfreezing conditions (Figures 1-2). This finding— diversification since the Southern Ocean reached its contemporary subfreezing temperature— generally aligns with findings for the Antarctic notothenioids (Near et al., 2015; Near et al., 2012). We saw less evidence for similar influence by Arctic Ocean cooling. A lack of influence by Arctic Ocean cooling on the evolutionary history of the Zoarcoidei could stem from the comparatively less harsh summer conditions of the Arctic versus Southern Ocean (e.g., water temperatures that are several degrees above zero, DeVries and Steffensen, 2005) reducing the ecological space for diversification (e.g., warmer water reducing the advantage of freezing tolerance), the more extreme physical isolation of the Southern Ocean relative to the Arctic Ocean, the more recent nature of Arctic cooling, or a combination of these, and perhaps other, factors.

In terms of topology, our phylogeny aligns with related efforts (Betancur-R et al., 2013; Kwun and Kim, 2013; Radchenko, 2016; Radchenko, 2015) and confirms standing taxonomic issues for the Zoarcoidei that have been noted previously (Kwun and Kim, 2013; Radchenko, 2016). We observed a lack of monophyly within the pricklebacks (family Stichaeidae). In some instances, taxa that are considered Stichaeidae are sister to other families (e.g., the Stichaeidae genus *Kasatkia* and Ptilichthyidae, posterior probability ≥ 0.95; Figures 1, S2), highlighting the need for the continued re-evaluation of higher-level taxonomic assignments within the suborder. Kwun and Kim (2013) addressed two of these issues by establishing two new families— Eulophiidae and Neozoarcidae—and reclassifying species previously considered to be Stichaeidae and Zoarcidae within them. Radchenko (2016) expanded on these descriptions, finding support for additional species to be grouped within both families. Our results support these taxonomic changes as well. Still, because Stichaeidae appear to have acted—at least in part—as a taxonomic “catch all” for the suborder, issues remain. For instance, *Poroclinus rothrocki* is currently assigned to Stichaeidae but we recovered it as sister to Cryptacanthodidae. Similarly, we recovered *Plectobranchus evides* (currently Stichaeidae) as sister to both *P. rothrocki* and *Zaprora silenus* (Zaproridae; Figure 1). Node probabilities for these three branches range from 0.61-0.85 (Figure S2) highlighting uncertainty around their placement. Thus, it is possible—and perhaps likely—that they each represent monotypic families similar to the prowfish (Zaproridae) but without additional analyses, ideally incorporating additional molecular data with morphological characters, it will remain uncertain. Finally, given monophyletic evidence in this study (posterior probability = 0.61; Figure S2) and the findings of Radchenko (2015), it may be warranted to elevate the subfamily Lumpeninae (Stichaeidae; Figure 1) to its own family, Lumpenidae.

### 4.2. Ancestral range estimations

Over 70 years ago, Shmidt (1950) hypothesized that major families in the suborder Zoarcoidei evolved in the northern Sea of Okhotsk (∼60°N) during the Miocene (23-5.5 Mya). In addition to our phylogenetic results supporting this timeline, our ancestral range reconstructions also supported it by showing that Zoarcoidei species largely diversified in mid-latitude regions of the Northern Hemisphere. In general, the ancestral range of a Zoarcoidei clade or lineage reflected its present-day distributions. This is particularly interesting in the context of eelpouts and their cosmopolitan distribution at both poles, the only family in the suborder to exhibit such a pattern (and one of only 10 families across all fishes, Møller et al., 2005). In addition to polar distributions, eelpouts are also the only Zoarcoidei family to commonly inhabit the deep sea (> 1000 m) and occur near hydrothermal vents (Møller et al., 2005). A wide range in preferred depths has been proposed as one factor that enhances geographical range size in marine organisms (Brown et al., 1996). This may be particularly true for deep water species like eelpouts given that the deep sea, while extreme in terms of pressure, cold, and darkness, is more environmentally consistent than shallower habitats and has few impediments to dispersal (Gaither et al., 2016). Thus, the global distribution of eelpouts relative to other families in the suborder (as well as their exceptionally high speciation rate, Rabosky et al., 2018) may be due to deep sea habitat connectivity paired with a propensity for adapting to extremes, whether subfreezing waters (Deng et al., 2010) or hydrothermal vents (Machida and Hashimoto, 2002).

### 4.3. Directionality of dispersal events

A striking biogeographic pattern within the Zoarcoidei is strong asymmetry in dispersal among geographic zones. For almost every pairing of geographic zones (e.g., Arctic and North Temperate), the rate of dispersal events was much higher from one zone into the other versus the reciprocal. This was most notable for the North Temperate zone, the center of origin for the group according to ancestral range reconstructions. Dispersals out of the North Temperate zone accounted for 30% of all events while dispersals into it only accounted for 5% (Figure 3A).

Similar patterns of asymmetric dispersal have been observed for other species, particularly from the North Pacific into the Arctic, for mollusks (Marincovich and Gladenkov, 1999) and other deep-water fishes (e.g., snailfishes, family Liparidae; Orr et al., 2019).

We also observed differences in dispersal rates for the Arctic and Antarctic. Given a relatively less harsh barrier to dispersal for species into or out of the Arctic versus Antarctic waters, which are surrounded by the Antarctic Circumpolar Current (ACC) and an extreme temperature drop (Barker et al., 2007), we expected more bidirectional dispersal for Arctic versus Antarctic. Our results, however, did not fully align with this expectation; while dispersal into the Arctic is indeed common (14% of all events), dispersal out of the Arctic is extremely rare (∼1%, Figure 3A). This starkly contrasts with the slightly higher but largely equivalent rates of dispersal into and out of the Antarctic (4% vs. 5% respectively, Figure 3A). Given the deep-water distributions of eelpouts and their tolerance for subfreezing temperatures, this result may be linked to differences in ecological opportunity or other factors between the regions. However, it might also simply reflect lineage age and species richness. The Arctic is adjacent to the North Temperate zone, the most likely center of origin for the group (and where much of its species richness remains), and by cooling much more recently, any barrier to dispersal that it presents are much younger than the Antarctic. Thus, a combination of geographic proximity to the Zoarcoidei center of origin paired with more recent thermal changes may be the most parsimonious explanation for the dispersal differences we observed between polar regions.

### 4.4. Potential caveats and future directions

Integrating phylogenetic insight with historical biogeographic modeling is a powerful approach for understanding the evolutionary history of organismal groups. When paired with well-studied environmental events (e.g., ocean cooling, continental separation), hypotheses about the relative importance of those events can be tested in a statistically robust framework. Still, this approach, and our implementation, is not without caveats that should be considered when interpreting our results and considering future studies.

The total numbers of biogeographic events reported in this study represent minima as we sampled ∼49% of the described species in the suborder. While more taxonomic sampling would provide greater resolution of the true value of these figures, it is unlikely to alter the relative proportions of each since, to our knowledge, no major bias in our sampling scheme exists in terms of both taxonomic representation and geographic scope. However, this only applies to the currently described taxonomic diversity. A more general, and important, caveat lies in the lack of knowledge surrounding Zoarcoidei species. Both eelpouts and the broader suborder are relatively deep-water taxa, living in hundreds to thousands of meters of water, with little biomedical or economic benefit. As such, they are understudied, and this lack of natural history knowledge may bias our results in two ways. First, many Zoarcoidei species have been described from the Sea of Okhotsk off the southeastern coast of Russia (∼55°N) and broadly from the Northern Hemisphere (Anderson, 1994). It is possible that a bias in both sampling effort and species descriptions towards the Northern Hemisphere, and specifically the North Temperate zone used in our study, influenced our results. However, our use of broad geographic zones likely tempered this effect as it allowed for broader distributions and therefore more uncertainty in species’ ranges. Second, most Zoarcoidei species have been described from morphology alone (Anderson, 1994) and little to no molecular insight exists for the group beyond phylogenies that target single representatives for each clade. Given the propensity for cryptic diversity even in well-studied groups (e.g., mouse lemurs, Hotaling et al., 2016) and the potential for morphologically distinct animals to be the same species (e.g., Jones and Weisrock, 2018), future efforts to assess species boundaries with molecular data across the suborder will improve resolution of their biogeographic history.

## 5. Conclusion

In this study, we used a densely sampled, time-calibrated phylogeny of the suborder Zoarcoidei, with an emphasis on the globally distributed eelpouts, to understand evolutionary relationships and biogeographic history for the group. From a taxonomic standpoint, we highlighted existing issues with the Zoarcoidei taxonomy and proposed new solutions. For biogeography, while our analyses at large geographic scales yielded key insights for the suborder and major clades, more targeted analyses of individual families paired with finer-scale distribution information and molecular data, will allow for testing more specific biogeographic hypotheses. Similarly, future efforts to use the same biogeographic methods across multiple taxonomic groups, perhaps comparing eelpouts to other deep, cold-water fauna (e.g., snailfishes), could shed additional light on how generalizable the role of major environmental changes like ocean cooling has been for fish diversification.

## 6. Acknowledgements

We acknowledge funding from the Antarctic Bursary, a Washington State University New Faculty Seed Grant, and NSF awards (OPP-1543383 and OPP-1947040 supporting T.D. and OPP-1906015 to J.L.K). We thank Keegan Paras and Charlotte Walker for their help with analyses. We also acknowledge the Computational Resources Core of the University of Idaho Institute for Bioinformatics and Evolutionary Studies (IBEST).

## 7. Data statement

The data (including sequence alignments), code, and additional results for this study are publicly available on Zenodo (https://doi.org/10.5281/zenodo.4306092).

## References

Altschul, S.F., Gish, W., Miller, W., Myers, E.W., Lipman, D.J., 1990. Basic local alignment search tool. Journal of Molecular Biology 215, 403–410. https://doi.org/10.1016/S0022-2836(05)80360-2

Anderson, M.E., 1994. Systematics and osteology of the Zoarcidae (Teleostei: Perciformes).

Ichthyological Bulletin of the J.L.B. Smith Institute of Ichthyology 60, 1–120.

Angelis, K., Álvarez-Carretero, S., Dos Reis, M., Yang, Z., 2018. An evaluation of different partitioning strategies for Bayesian estimation of species divergence times. Systematic Biology 67, 61–77. https://doi.org/10.1093/sysbio/syx061

Bankevich, A., Nurk, S., Antipov, D., Gurevich, A.A., Dvorkin, M., Kulikov, A.S., Lesin, V.M., Nikolenko, S.I., Pham, S., Prjibelski, A.D., 2012. SPAdes: a new genome assembly algorithm and its applications to single-cell sequencing. Journal of Computational Biology 19, 455–477. https://doi.org/10.1089/cmb.2012.0021

Barker, P.F., Filippelli, G.M., Florindo, F., Martin, E.E., Scher, H.D., 2007. Onset and role of the Antarctic Circumpolar Current. Deep Sea Research Part II: Topical Studies in Oceanography 54, 2388–2398. https://doi.org/10.1016/j.dsr2.2007.07.028

Betancur-R, R., Broughton, R.E., Wiley, E.O., Carpenter, K., López, J.A., Li, C., Holcroft, N.I., Arcila, D., Sanciangco, M., Cureton Ii, J.C., 2013. The tree of life and a new classification of bony fishes. PLoS currents 5. https://doi.org/10.1371/currents.tol.53ba26640df0ccaee75bb165c8c26288

Bieńkowska-Wasiluk, M., Bonde, N., Møller, P.R., Gaździcki, A., 2013. Eocene relatives of cod icefishes (perciformes: Notothenioidei) from Seymour Island, Antarctica. Geological Quarterly 57, 567–582, doi: 510.7306/gq.1112. https://doi.org/10.7306/gq.1112

Brown, J.H., Stevens, G.C., Kaufman, D.M., 1996. The geographic range: size, shape, boundaries, and internal structure. Annual Review of Ecology and Systematics 27, 597–623. https://doi.org/10.1146/annurev.ecolsys.27.1.597

Cavanaugh, J.E., 1997. Unifying the derivations for the Akaike and corrected Akaike information criteria. Statistics & Probability Letters 33, 201–208. https://doi.org/10.1016/S0167-7152(96)00128-9

Chen, L., DeVries, A.L., Cheng, C.H., 1997. Evolution of antifreeze glycoprotein gene from a trypsinogen gene in Antarctic notothenioid fish. Proceedings of the National Academy of Sciences 94, 3811–3816. https://doi.org/10.1073/pnas.94.8.3811

Chernomor, O., Von Haeseler, A., Minh, B.Q., 2016. Terrace aware data structure for phylogenomic inference from supermatrices. Systematic Biology 65, 997–1008. https://doi.org/10.1093/sysbio/syw037

Clardy, T.R., 2014. Phylogenetic systematics of the prickleback family Stichaeidae (Cottiformes: Zoarcoidei) using morphological data. Dissertations, Theses, and Masters Projects. 1539616612. https://doi.org/10.25773/v5-fyer-5n47

Davies, P.L., Baardsnes, J., Kuiper, M.J., Walker, V.K., 2002. Structure and function of antifreeze proteins. Philosophical transactions of the Royal Society of London. Series B, Biological sciences 357, 927–935. https://doi.org/10.1098/rstb.2002.1081

Davies, P.L., Hew, C.L., Fletcher, G.L., 1988. Fish antifreeze proteins: physiology and evolutionary biology. Canadian Journal of Zoology 66, 2611–2617. https://doi.org/10.1139/z88-385

Deng, C., Cheng, C.H., Ye, H., He, X., Chen, L., 2010. Evolution of an antifreeze protein by neofunctionalization under escape from adaptive conflict. Proceedings of the National Academy of Sciences of the United States of America 107, 21593–21598. https://doi.org/10.1073/pnas.1007883107

DeVries, A.L., Steffensen, J.F., 2005. The Arctic and Antarctic polar marine environments. Fish physiology 22, 1–24. https://doi.org/10.1016/S1546-5098(04)22001-5

Dupin, J., Matzke, N.J., Särkinen, T., Knapp, S., Olmstead, R.G., Bohs, L., Smith, S.D., 2017. Bayesian estimation of the global biogeographical history of the Solanaceae. Journal of Biogeography 44, 887–899. https://doi.org/10.1111/jbi.12898

Edgar, R.C., 2004. MUSCLE: multiple sequence alignment with high accuracy and high throughput. Nucleic acids research 32, 1792–1797. https://doi.org/10.1093/nar/gkh340

Fitch, J., 1967. The marine fish fauna, based primarily on otoliths, of a lower Pleistocene deposit at San Pedro, California (LACMIP 332, San Pedro Sand). Los Angeles County Museum of Natural History.

Fricke, R., Eschmeyer, W., Van der Laan, R., 2018. Catalog of fishes: genera, species, references. California Academy of Sciences, San Francisco, CA, USA

Froese, R., Pauly, D., 2019. FishBase in the Catalogue of Life.

Gaither, M.R., Bowen, B.W., Rocha, L.A., Briggs, J.C., 2016. Fishes that rule the world: circumtropical distributions revisited. Fish and Fisheries 17, 664–679. https://doi.org/10.1111/faf.12136

González-Wevar, C.A., Nakano, T., Cañete, J.I., Poulin, E., 2010. Molecular phylogeny and historical biogeography of Nacella (Patellogastropoda: Nacellidae) in the Southern Ocean. Molecular phylogenetics and evolution 56, 115–124. https://doi.org/10.1016/j.ympev.2010.02.001

Grassle, J.F., 2000. The Ocean Biogeographic Information System (OBIS): an on-line, worldwide atlas for accessing, modeling and mapping marine biological data in a multidimensional geographic context. Oceanography 13, 5–7.

Heath, T.A., Huelsenbeck, J.P., Stadler, T., 2014. The fossilized birth–death process for coherent calibration of divergence-time estimates. Proceedings of the National Academy of Sciences 111, E2957–E2966. https://doi.org/10.1073/pnas.1319091111

Hoang, D.T., Chernomor, O., Von Haeseler, A., Minh, B.Q., Vinh, L.S., 2018. UFBoot2: improving the ultrafast bootstrap approximation. Molecular biology and evolution 35, 518–522. https://doi.org/10.1093/molbev/msx281

Hopkins, D., Marincovich Jr, L., 1984. Whale biogeography and the history of the Arctic Basin. Works of the Arctic Centre 8, 7–24.

Hotaling, S., Foley, M.E., Lawrence, N.M., Bocanegra, J., Blanco, M.B., Rasoloarison, R., Kappeler, P.M., Barrett, M.A., Yoder, A.D., Weisrock, D.W., 2016. Species discovery and validation in a cryptic radiation of endangered primates: coalescent-based species delimitation in Madagascar’s mouse lemurs. Molecular ecology 25, 2029–2045. https://doi.org/10.1111/mec.13604

Jones, K.S., Weisrock, D.W., 2018. Genomic data reject the hypothesis of sympatric ecological speciation in a clade of Desmognathus salamanders. Evolution 72, 2378–2393. https://doi.org/10.1111/evo.13606

Kalyaanamoorthy, S., Minh, B.Q., Wong, T.K., von Haeseler, A., Jermiin, L.S., 2017. ModelFinder: fast model selection for accurate phylogenetic estimates. Nature Methods 14, 587–589. https://doi.org/10.1038/nmeth.4285

Kwun, H.J., Kim, J.-K., 2013. Molecular phylogeny and new classification of the genera Eulophias and Zoarchias (PISCES, Zoarcoidei). Molecular Phylogenetics and Evolution 69, 787–795. https://doi.org/10.1016/j.ympev.2013.06.025

Landis, M.J., Matzke, N.J., Moore, B.R., Huelsenbeck, J.P., 2013. Bayesian analysis of biogeography when the number of areas is large. Systematic Biology 62, 789–804. https://doi.org/10.1093/sysbio/syt040

Lane, M.A., Edwards, J.L., 2007. The global biodiversity information facility (GBIF). Biodiversity databases: Techniques, politics, and applications, 1–4.

Machida, Y., Hashimoto, J., 2002. Pyrolycus manusanus, a new genus and species of deep-sea eelpout from a hydrothermal vent field in the Manus Basin, Papua New Guinea (Zoarcidae, Lycodinae). Ichthyological Research 49, 1–6. https://doi.org/10.1007/s102280200000

Marincovich, L., Gladenkov, A.Y., 1999. Evidence for an early opening of the Bering Strait. Nature 397, 149–151. https://doi.org/10.1038/16446

Matschiner, M., Hanel, R., Salzburger, W., 2011. On the origin and trigger of the notothenioid adaptive radiation. PloS one 6, e18911. https://doi.org/10.1371/journal.pone.0018911

Matzke, N.J., 2013. Probabilistic historical biogeography: new models for founder-event speciation, imperfect detection, and fossils allow improved accuracy and model-testing. Frontiers of Biogeography 5. https://doi.org/10.21425/F5FBG19694

Matzke, N.J., 2014. Model selection in historical biogeography reveals that founder-event speciation is a crucial process in island clades. Systematic Biology 63, 951–970. https://doi.org/10.1093/sysbio/syu056

Møller, P.R., Nielsen, J.G., Anderson, M.E., 2005. Systematics of polar fishes. Fish Physiology 22, 25–78. https://doi.org/10.1016/S1546-5098(04)22002-7

Nauheimer, L., Metzler, D., Renner, S.S., 2012. Global history of the ancient monocot family Araceae inferred with models accounting for past continental positions and previous ranges based on fossils. New Phytologist 195, 938–950. https://doi.org/10.1111/j.1469-8137.2012.04220.x

Nazarkin, M., 1998. New stichaeid fishes (Stichaeidae, Perciformes) from Miocene of Sakhalin. Journal of Ichthyology 38, 279–291.

Nazarkin, M., 2002. Gunnels (Perciformes, Pholidae) from the Miocene of Sakhaline Island. Journal of Ichthyology 42, 279–288.

Nazarkin, M., Yabumoto, Y., 2015. New fossils of Neogene pricklebacks (Actinopterygii: Stichaeidae) from East Asia. Zoosystematica Rossica 24, 128–137.

Near, T.J., Dornburg, A., Harrington, R.C., Oliveira, C., Pietsch, T.W., Thacker, C.E., Satoh, T.P., Katayama, E., Wainwright, P.C., Eastman, J.T., 2015. Identification of the notothenioid sister lineage illuminates the biogeographic history of an Antarctic adaptive radiation. BMC evolutionary biology 15, 109. https://doi.org/10.1186/s12862-015-0362-9

Near, T.J., Dornburg, A., Kuhn, K.L., Eastman, J.T., Pennington, J.N., Patarnello, T., Zane, L., Fernández, D.A., Jones, C.D., 2012. Ancient climate change, antifreeze, and the evolutionary diversification of Antarctic fishes. Proceedings of the National Academy of Sciences 109, 3434–3439. https://doi.org/10.1073/pnas.1115169109

Nguyen, L.-T., Schmidt, H.A., Von Haeseler, A., Minh, B.Q., 2015. IQ-TREE: a fast and effective stochastic algorithm for estimating maximum-likelihood phylogenies. Molecular biology and evolution 32, 268–274. https://doi.org/10.1093/molbev/msu300

Orr, J.W., Spies, I., Stevenson, D.E., Longo, G.C., Kai, Y., Ghods, S., Hollowed, M., 2019. Molecular phylogenetics of snailfishes (Cottoidei: Liparidae) based on MtDNA and RADseq genomic analyses, with comments on selected morphological characters. Zootaxa 4642, 1–79. https://doi.org/10.1016/j.ympev.2004.06.015

Paradis, E., Schliep, K., 2019. ape 5.0: an environment for modern phylogenetics and evolutionary analyses in R. Bioinformatics 35, 526–528. https://doi.org/10.1093/bioinformatics/bty633

Quinlan, A.R., Hall, I.M., 2010. BEDTools: a flexible suite of utilities for comparing genomic features. Bioinformatics 26, 841–842. https://doi.org/10.1093/bioinformatics/btq033

Rabosky, D.L., Chang, J., Title, P.O., Cowman, P.F., Sallan, L., Friedman, M., Kaschner, K., Garilao, C., Near, T.J., Coll, M., 2018. An inverse latitudinal gradient in speciation rate for marine fishes. Nature 559, 392–395. https://doi.org/10.1038/s41586-018-0273-1

Radchenko, O., 2016. Timeline of the evolution of eelpouts from the suborder Zoarcoidei (Perciformes) based on DNA variability. Journal of Ichthyology 56, 556–568. https://doi.org/10.1134/S0032945216040123

Radchenko, O.A., 2015. The System of the Suborder Zoarcoidei (Pisces, Perciformes) as Inferred from Molecular Genetic Data. Russian Journal of Genetics 51, 1273–1290. https://doi.org/10.1134/S1022795415100130

Ree, R.H., 2005. Detecting the historical signature of key innovations using stochastic models of character evolution and cladogenesis. Evolution 59, 257–265. https://doi.org/10.1111/j.0014-3820.2005.tb00986.x

Ree, R.H., Smith, S.A., 2008. Maximum likelihood inference of geographic range evolution by dispersal, local extinction, and cladogenesis. Systematic Biology 57, 4–14. https://doi.org/10.1080/10635150701883881

Revell, L.J., 2012. phytools: an R package for phylogenetic comparative biology (and other things). Methods in Ecology and Evolution 3, 217–223. https://doi.org/10.1111/j.2041-210X.2011.00169.x

Ronquist, F., 1997. Dispersal-vicariance analysis: a new approach to the quantification of historical biogeography. Systematic Biology 46, 195–203. https://doi.org/10.1093/sysbio/46.1.195

Ronquist, F., Teslenko, M., Van Der Mark, P., Ayres, D.L., Darling, A., Höhna, S., Larget, B., Liu, L., Suchard, M.A., Huelsenbeck, J.P., 2012. MrBayes 3.2: efficient Bayesian phylogenetic inference and model choice across a large model space. Systematic biology 61, 539–542. https://doi.org/10.1093/sysbio/sys029

Schluter, D., 2000. The ecology of adaptive radiation. OUP Oxford.

Shmidt, P., 1950. Ryby Okhotskogo Morya [Fishes of the Sea of Okhotsk]. Trudy Tikhookeanskogo Komiteta 6, 1–370.

Suyama, M., Torrents, D., Bork, P., 2006. PAL2NAL: robust conversion of protein sequence alignments into the corresponding codon alignments. Nucleic acids research 34, W609–W612. https://doi.org/10.1093/nar/gkl315

Yabumoto, Y., Uyeno, T., 1994. Late Mesozoic and Cenozoic fish faunas of Japan. Island Arc 3, 255–269. https://doi.org/10.1111/j.1440-1738.1994.tb00115.x

Zhang, C., Stadler, T., Klopfstein, S., Heath, T.A., Ronquist, F., 2016. Total-evidence dating under the fossilized birth–death process. Systematic Biology 65, 228–249. https://doi.org/10.1093/sysbio/syv080

