## Supplementary Figures S1-S3 for "The biogeographic history of eelpouts and related fishes: linking phylogeny, environmental change, and patterns of dispersal in a globally distributed fish group"

**Figure S1.** A dated Bayesian phylogeny of the suborder Zoarcoidei. A simplified version is included as Figure 1.

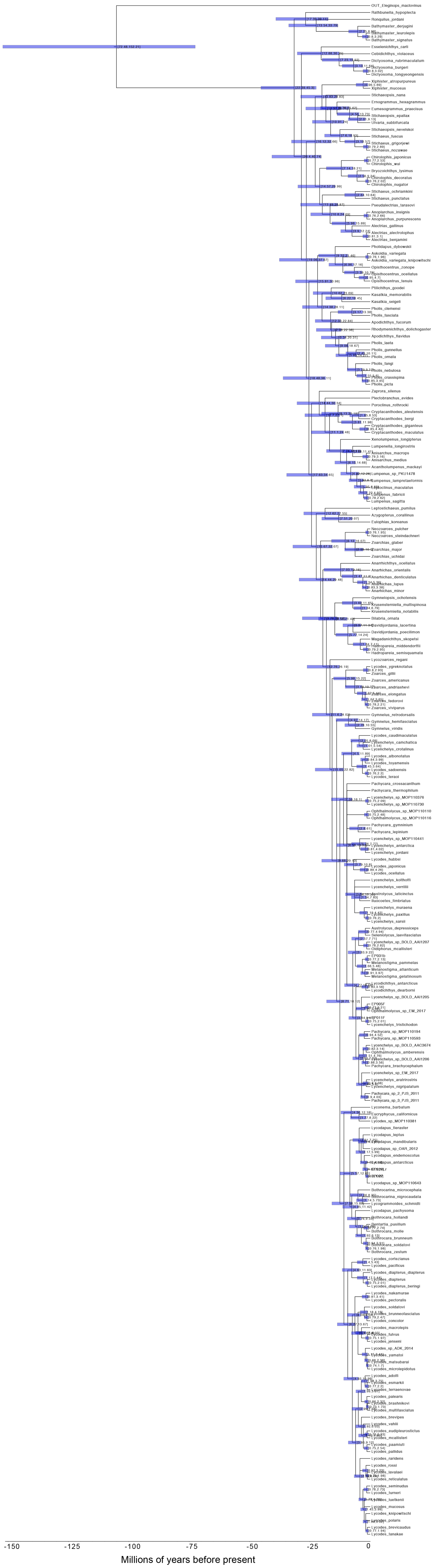

**Figure S2.** A dated Bayesian phylogeny of the suborder Zoarcoidei with node probabilities shown. A simplified version is included as Figure 1.

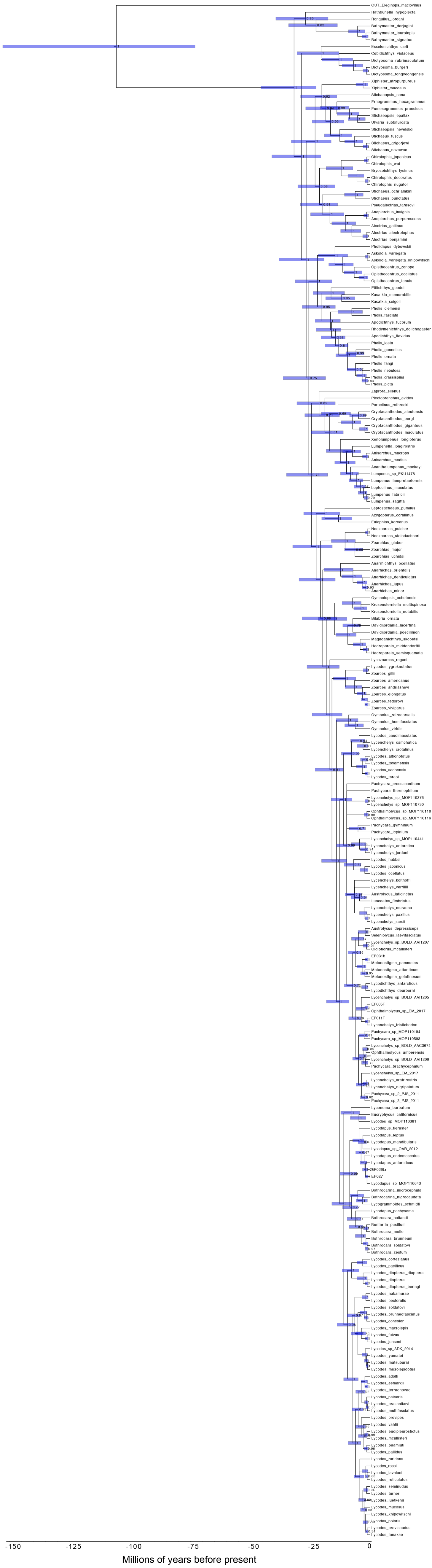

**Figure S3.** The full maximum-likelihood phylogeny of the suborder Zoarcoidei based on an IQ-TREE analysis.

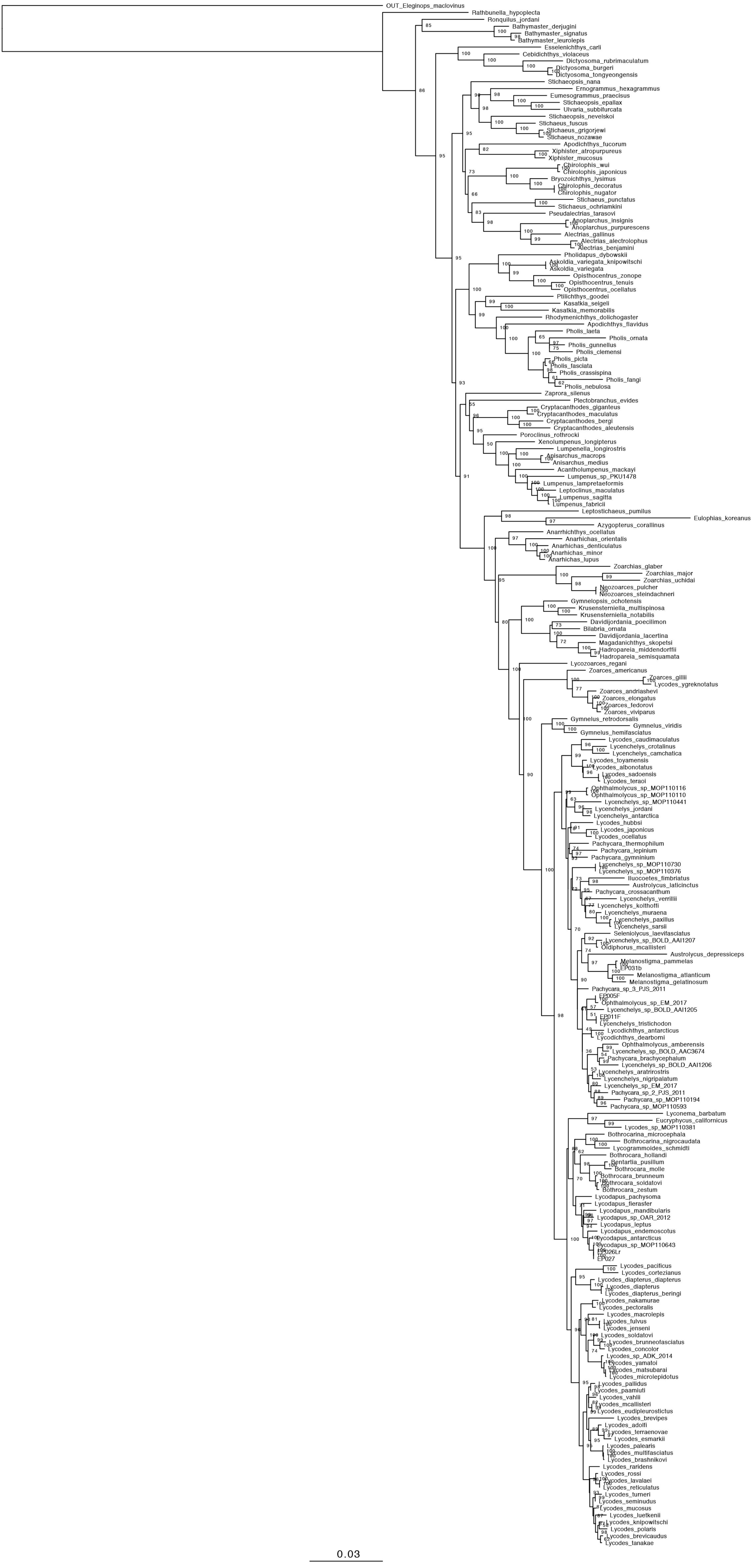
